# Repetitive DNA shapes genome architecture and chromosomal diversification in birds of prey

**DOI:** 10.64898/2026.03.03.709396

**Authors:** Guilherme Mota Souza, Jhon Alex Dziechciarz Vidal, Gustavo Akira Toma, Rafael Kretschmer, Edivaldo Herculano Correa de Oliveira, Thomas Liehr, Marcelo de Bello Cioffi

**Affiliations:** Departamento de Genética e Evolução, Universidade Federal de São Carlos, 13565-905 São Carlos, SP, Brasil; Laboratório de Citogenética e Evolução, Departamento de Genética, Instituto de Biociências, Universidade Federal do Rio Grande do Sul, Porto Alegre 91509-900, RS, Brazil; Seção de Meio Ambiente, Instituto Evandro Chagas, 67030-000 Ananindeua, PA, Brazil; Instituto de Ciências Exatas e Naturais, Universidade Federal do Pará, 66075-110 Belém, PA, Brazil; Jena University Hospital, Friedrich Schiller University, Institute of Human Genetics, Jena, Germany

**Keywords:** Accipitridae, Repeatome, Cytogenomics, Genome architecture dynamics

## Abstract

The evolution of genome architecture occurs through dynamic interactions between repetitive DNAs and chromosomal organization; nevertheless, the processes underlying these mechanisms are not well understood. This study presents a comprehensive genomic and cytogenetic analysis of repetitive DNA evolution across Accipitridae birds, a raptor family notable for its significant chromosomal variation. We aimed to investigate how repetitive DNAs have evolved across Accipitriform lineages and test whether shifts in repeat composition are associated with patterns of species diversification. Comparative investigations of eight genomes reveal lineage-specific spikes of transposable elements and satellite DNAs that substantially modify genome composition while preserving a common structural framework. Temporal insertion profiles indicate that repeat turnover is ongoing and frequently coincides with lineages exhibiting extensive chromosomal reorganization. By integrating comparative repeatome analyses with *in silico* and cytogenetic mapping, we elucidate the spatial architecture governing repeat dynamics, connecting molecular turnover to their chromosomal structure. These findings underscore the effectiveness of merging genomic and chromosomal data to elucidate the impact of repeat landscapes on chromosomal and genomic evolution.

## Introduction

Genome architecture evolves through a dynamic interplay between large-scale structural rearrangements and the expansion or contraction of repetitive DNAs, processes that collectively modify the chromosomal organization and gene landscapes over the course of evolution^1-3^. Increasing evidence indicates that repetitive elements, including transposable elements and satellite DNAs, are key contributors to chromosomal remodeling^4-7^, yet their comprehensive evolutionary impact on genome architecture remains poorly understood. Lineage-specific bursts of repeat amplification can influence recombination landscapes, heterochromatin formation, and chromosome stability, with downstream consequences for karyotype diversification^3^. Comparative genomic studies have revealed that repeat landscapes can differ markedly even among closely related taxa, raising the possibility that repeat turnover may be linked to evolutionary trajectories^8^. Notwithstanding these findings, the degree to which repeat dynamics correlate with genomic architecture and diversification patterns, especially in groups exhibiting significant karyotypic variability, remains largely unexamined.

Birds are notable for their longstanding structural stability, with significantly fewer large-scale chromosomal rearrangements than most other vertebrate lineages^9-12^. This karyotypic conservatism has been interpreted as a defining feature of avian genome evolution, reflecting strong constraints on chromosome organization over deep evolutionary timescales. This stability, however, is not consistent among all bird lineages. Several clades exhibit significant deviations from the ancestral pattern (2n=80), demonstrating considerable chromosomal reshuffling and structural reorganization (reviewed in ^13^). These lineage-specific bursts of rearrangement underscore that avian genome evolution is influenced by a balance between long-term conservation and intermittent structural alteration. Accipitriformes, which includes eagles, harriers, hawks, kites, and erne, represents one such lineage, exhibiting extensive chromosomal reorganization. This reorganization has been driven not only by recurrent fusions between macro- and microchromosomes but also by multiple fissions involving the largest ancestral chromosomal pairs, along with marked variation in diploid chromosome numbers (2n)^12,14-16^. Within this order, the family Accipitridae, encompassing more than 250 extant species and representing the largest radiation of diurnal birds of prey^17^, exhibits a particularly striking chromosomal diversity. Besides the predominant 2n of 66 in Accipitrids, extreme cases of chromosomal reduction are observed, such as 2n = 58 in *Harpia harpyja*^18,19^ and 2n = 54 in *Morphnus guianensis*^20^, underscoring the remarkable karyotypic diversity of the lineage. Owing to their diversity, evolutionary breadth, and pronounced structural variation, accipitrids provide a powerful comparative framework for investigating the processes that shape avian genome architecture.

Recent advancements in genome sequencing have enhanced the potential to characterize the whole set of repetitive DNAs (Repeatome) and their evolutionary dynamics among species. Specifically, repeatome profiling with low-coverage sequencing has emerged as an efficient and cost-effective method for characterizing the main categories of repeated elements and comparing genomic compositions across taxa, allowing the identification of lineage-specific expansions and contractions^21,22^. The increasing availability of chromosome-level genome assemblies opens up the interpretation of these repeat landscapes within a chromosomal framework, correlating variations in repeat composition with genome size, chromosomal architecture, and genome reorganization patterns^8,23^. When combined with time-calibrated phylogenies, such genomic datasets provide an opportunity to test whether shifts in repetitive DNA content correlate with broader evolutionary processes, including chromosomal evolution and lineage diversification^24,25^. Cytogenetic analyses, especially the *in situ* mapping of repetitive elements, enhance sequence-based methods by displaying their physical chromosomal distribution, thereby integrating molecular and karyotypic perspectives and providing a more thorough understanding of repeat landscape evolution across species^26^. Here, we investigate the evolution of repetitive DNAs across eight accipitrid birds through an integrative framework that combines comparative genomics with targeted cytogenetic analysis. These selected species present a marked chromosomal diversity **(Supplementary Figure 1)**, also reflected by a variation in their genome sizes **(Figure 1)**. Broad-scale *in silico* comparisons of available accipitrid genomes were used to contextualize repeat landscape variation in an evolutionary framework, whereas empirical repeatome profiling and cytogenetic validation were performed in the harpy eagle (*Harpia harpyja*) as a focal model species. Using low-coverage sequencing, insertion age analyses, and both *in silico* and *in situ* localization of repetitive elements, we evaluated how specific repeat classes contribute to genome organization. Comparative analyses enabled us to distinguish lineage-associated repeat signatures from more conserved patterns, while the detailed cytogenetic characterization of *H. harpyja* provided chromosomal resolution linking sequence composition to structural architecture. This approach provided a multi-scale viewpoint on the potential impact of repetitive DNAs on genome evolution and chromosomal reorganization in birds of prey.

**Figure 1.**
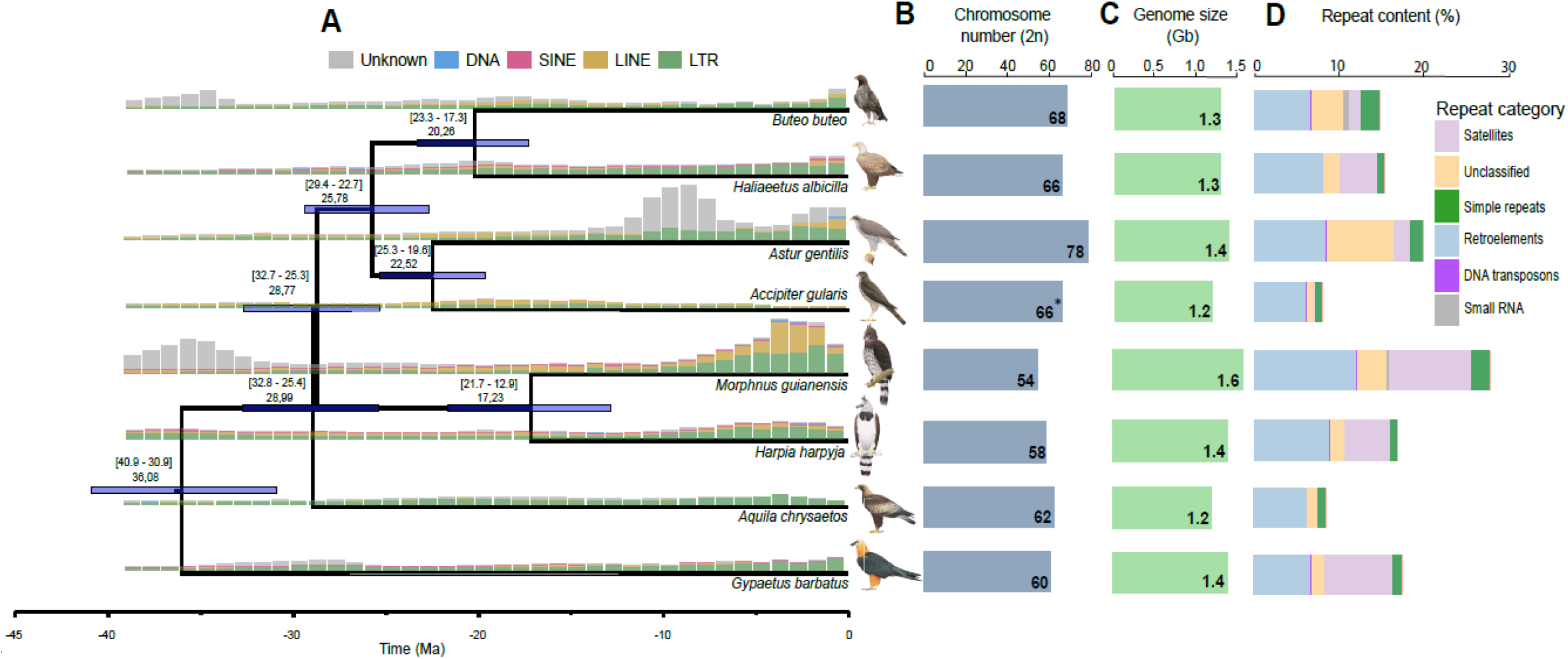
Phylogenetic and comparative analyses of eight Accipitridae genomes. (A) Phylogeny of the eight Accipitridae species, showing their divergence times according to^27^. The mean divergence time is labeled at each node. The vertical bars show the frequency of TE insertions during the evolution of these species (see details in “Methods”). (B) The diploid numbers of each species are displayed, along with their respective genome sizes (C) and repeat content (D). Asterisks denote the estimated diploid number for *Accipiter gularis* based on Hi-C scaffolding, without cytogenetic evidence.

## Results

### Characterization and comparison of Accipitridae repeatomes

A comparative investigation of Accipitridae revealed that most species experienced distinct patterns of repetitive DNA accumulation, except *A. chrysaetus* and *A. gularis*, which showed similar proportions of repeat classes **(Figure 1; Supplementary Figure 2)**. Differential amplification of three major repeat classes contributed to the interspecific variation observed in their repetitive DNA content. Retroelements were the dominant repeat category across all eight species. Satellite DNAs were the second most abundant class in *G. barbatus, H. harpyja*, and *H. albicilla*, whereas *B. buteo* differed from the other species by exhibiting additional amplification of small RNA elements **(Supplementary Table 2)**. A refined comparison of relative retroelement abundance **(Supplementary Figure 3)** revealed that LTR retrotransposons, retroviral elements, LINEs, and L2/CR1/Rex consistently represented the most abundant retroelement families across Accipitridae. In contrast, other families showed lineage-specific amplification patterns, with some absent from particular species, including the R2/R4/NeSL group.

The temporal landscapes of repetitive DNA accumulation revealed interspecific differences in repeat amplification **(Figure 1A)**. In all species, LTR elements dominated the repetitive fraction, which is expected, since they represent the most abundant repeat class in all genomes. However, the timing and intensity of amplification events varied among species.

*M. guianensis* and *A. gentilis* exhibited pronounced medium-to-recent bursts of repetitive DNA accumulation, particularly driven by LINEs and LTR elements. Divergence time estimates indicate that major amplification pulses occurred predominantly after the divergence of major accipitrid lineages, highlighting independent and species-specific trajectories of repetitive DNA evolution **(Figure 1)**. K2P divergence profiles **(Supplementary Figure 2)** further indicated that species with higher SatDNA abundance experienced relatively recent to intermediate amplification of this repeat class, suggesting that the expansion of SatDNAs is a comparatively recent feature of Accipitridae genomes. Genome sizes ranged from ∼1.2 to 1.6 Gb and covaried with total repeat content, while diploid chromosome numbers differed substantially among lineages **(Figure 1)**. Species with reduced chromosome counts showed distinct repeat class distributions relative to taxa with higher counts, indicating that repeatomes share a common structural framework but vary quantitatively among species.

### Chromosome repeat landscapes in Accipitridae

Across the eight Accipitridae genomes, repetitive DNAs displayed a species-specific chromosomal distribution pattern, forming prominent high-density clusters within a background of lower-density dispersion. Although repeats are preferentially enriched in terminal and pericentromeric regions in a broadly conserved genomic architecture, the extent and intensity of these accumulations vary markedly among species **(Figure 2)**. Several macrochromosomes in *G. barbatus, M. guianensis*, and *H. harpyja* contain prominent repeat-rich blocks, consistent with lineage-specific amplification or retention of repetitive sequences. By contrast, *A. gentilis* and *A. gularis* display more dispersed and fragmented patterns, suggesting alternative modes of repeat organization **(Figure 2)**.

**Figure 2.**
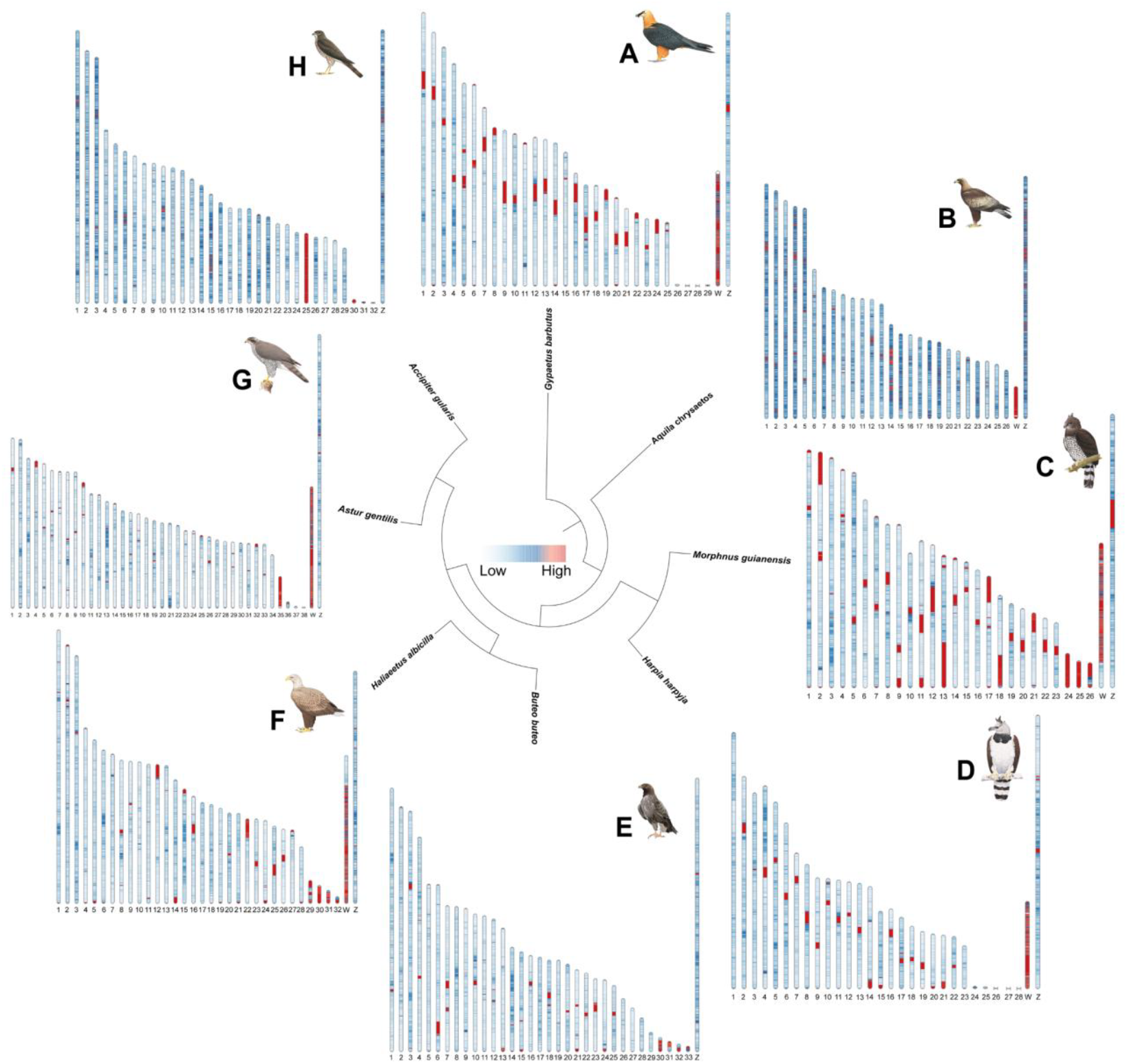
Chromosomal density of repetitive DNA across A, *Gypaetus barbatus*; B, *Aquila chrysaetos*; C, *Morphnus guianensis;* D, *Harpia harpyja;* E, *Buteo buteo;* F, *Haliaeetus albicilla;* G, *Astur gentilis*, and H, *Accipiter gularis* chromosomes.

Although the sequenced genome of *A. gularis* originates from a male individual, its chromosomal repeat landscape suggests that chromosome 25 likely corresponds to the W chromosome **(Figure 2H)**. This interpretation is supported by the markedly elevated repetitive DNA density of this chromosome, which is similar to the repeat enrichment seen on the W chromosome in the other species **(Figure 2)**. Consistently, synteny analyses indicate homology between chromosome 25 and the W chromosome of other taxa **(Supplementary Figure 1)**

### General features of Accipitridae satellitomes

Satelliteome characterization identified 18 SatDNA families for *A. chrysaetos* (AchSat) and *A. gularis* (AguSat), 16 for *A. gentilis* (AgeSat) and *B. buteo* (BbuSat), 15 for *H. harpyja* (HhaSat), and only 8 families for *H. albicilla* **(Tables Supplementary 3-8)**. Among the isolated SatDNAs, the majority exhibited an elevated GC content. Specifically, GC-rich satellites accounted for 66.6% in *A. chrysaetos*, 73.3% in *H. harpyja*, 72.2% in *A. gularis*, 81.2% in *A. gentilis*, 87.5% in *H. albicilla*, and 56.2% in *B. buteo*. Overall, approximately 70% of the isolated SatDNAs displayed higher GC content. The lengths of the repeating units (RULs) varied among species, ranging from 27 bp (AchSat15) to 4,231 bp (AchSat05) in *A* .*chrysaetos*; from 29 bp (AguSat10) to 1,779 bp (AguSat07) in *A. gularis*; from 32 bp (AgeSat08 and AgeSat15) to 7,330 bp in *A. gentilis*; from 27 bp (BbuSat10) to 2,179 bp (BbuSat08) in *B. buteo*; from 31 bp (HhaSat13) to 2,584 bp (HhaSat06) in *H. harpyja*; and from 33 bp (HalSat03) to 1,386 bp (HalSat06) in *H. albicilla*. Overall, approximately 70% of the SatDNA families exhibited RULs greater than 150 bp (**Tables Supplementary 3-8**).

We identified 8 superfamilies shared among the species, encompassing approximately 75% of all SatDNA families (**Supplementary Table 9**). Next, alignment and homology analyses were used to identify and isolate, within each group, only the variants shared between species **(Table 1)**.

**Table 1.** SatDNA variants identified within each superfamily.

| Group | *Harpia harpyja* | *Accipiter gularis* | *Astur gentilis* | *Aquila chrysaetos* | *Buteo buteo* | *Haliaeetus albicilla* | Similarity (%) |
| --- | --- | --- | --- | --- | --- | --- | --- |
| Group 2 | HhaSat01-173 | AguSat04-173 | AgeSat01-173 | AchSat01-173 | BbuSat01-173 | HalSat01-173 | 78.5 - 93.1 |
|  |  |  |  | AchSat15-27 | BbuSat10-27 |  | 92.9 |
|  | HhaSat09-63 |  |  | AchSat14-65 |  |  | 93.8 |
| Group 3 | HhaSat03-33 | AguSat15-36 | AgeSat08-32 |  | BbuSat09-32 |  | 76 - 84.8 |
|  |  | AguSat13-376 | AgeSat02-595 |  |  |  | 82.8 |
|  |  |  |  |  | BbuSat05-379 | HalSat02-444 | 87 |
| Group 4 |  |  |  | AchSat09-2148 | BbuSat08-2179 |  | 82.4 - 86.8 |
| Group 5 | HhaSat12-539 |  | AgeSat16-534 | AchSat08-537 | BbuSat12-539 | HalSat05-539 | 86.1 - 92.4 |
| Group 6 | HhaSat14-1373 |  | AgeSat13-1389 | AchSat10-1215 | BbuSat07-1374 | HalSat06-1386 | 80.1 - 96.1 |
| Group 8 |  | AguSat06-343 | AgeSat07-342 |  | BbuSat06-342 |  | 89 - 94.5 |

### Chromosomal architecture of shared SatDNAs validated by in silico and in situ mapping

The *in silico* chromosomal mapping demonstrates that shared SatDNA families display a notable distribution pattern throughout the chromosomes of all eight accipitrid species, with recurrent clustering in pericentromeric and subtelomeric areas and frequent enrichment on sex chromosomes **(Figure 3)**. Several SatDNA groups are preserved across phylogenetically distant species, occurring on homologous macrochromosomes and subsets of microchromosomes, which implies long-term preservation of ancestral repetitions. This conservation is particularly noticeable in SatDNA families that are present on large autosomes and on the Z chromosome. At the same time, each species shows a unique pattern, showing lineage-specific amplification or contraction events **(Figure 3)**.

**Figure 3.**
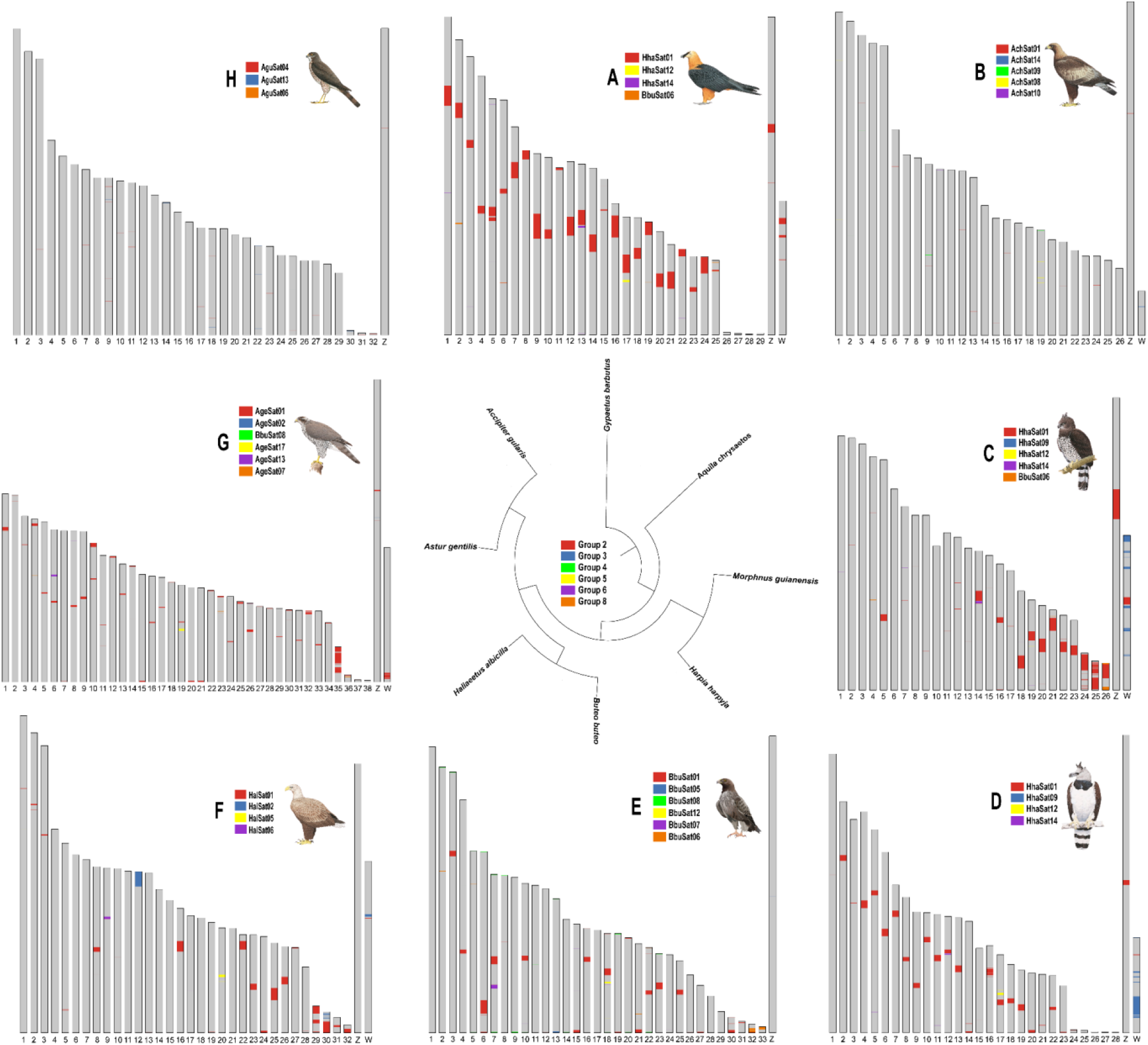
*In silico* chromosomal mapping of shared satellite DNA families in A, *Gypaetus barbatus*; B, *Aquila chrysaetos*; C, *Morphnus guianensis;* D, *Harpia harpyja;* E, *Buteo buteo;* F, *Haliaeetus albicilla;* G, *Astur gentilis*, and H, *Accipiter gularis*. Colors denote individual SatDNA families, as indicated in the central legend.

Due to the availability of chromosomal preparations solely for *Harpia harpyja*, we conducted *in situ* mapping of its SatDNA families to empirically confirm the chromosomal distribution predicted from the *in silico* analysis **(Figure 3D)**. The *in situ* chromosomal mapping confirms the primary distribution patterns anticipated by the *in silico* analysis, demonstrating a significant consistency in both chromosomal localization and relative signal enrichment **(Figure 4)**. HhaSat01, HhaSat02, and HhaSat06 were dispersed across multiple chromosomes, although absent from specific autosomal pairs and the ZW sex chromosomes. In contrast, HhaSat03, HhaSat12, and HhaSat13 showed progressively restricted localization to two or a single autosomal pair. HhaSat10 and HhaSat11 displayed strict sex-chromosome specificity, being located exclusively on the Z and W chromosomes, respectively. HhaSat15 was confined to a telomeric region of a single chromosome, whereas HhaSat04 and HhaSat14 produced no detectable signals under the conditions tested **(Figure 4)**.

**Figure 4.**
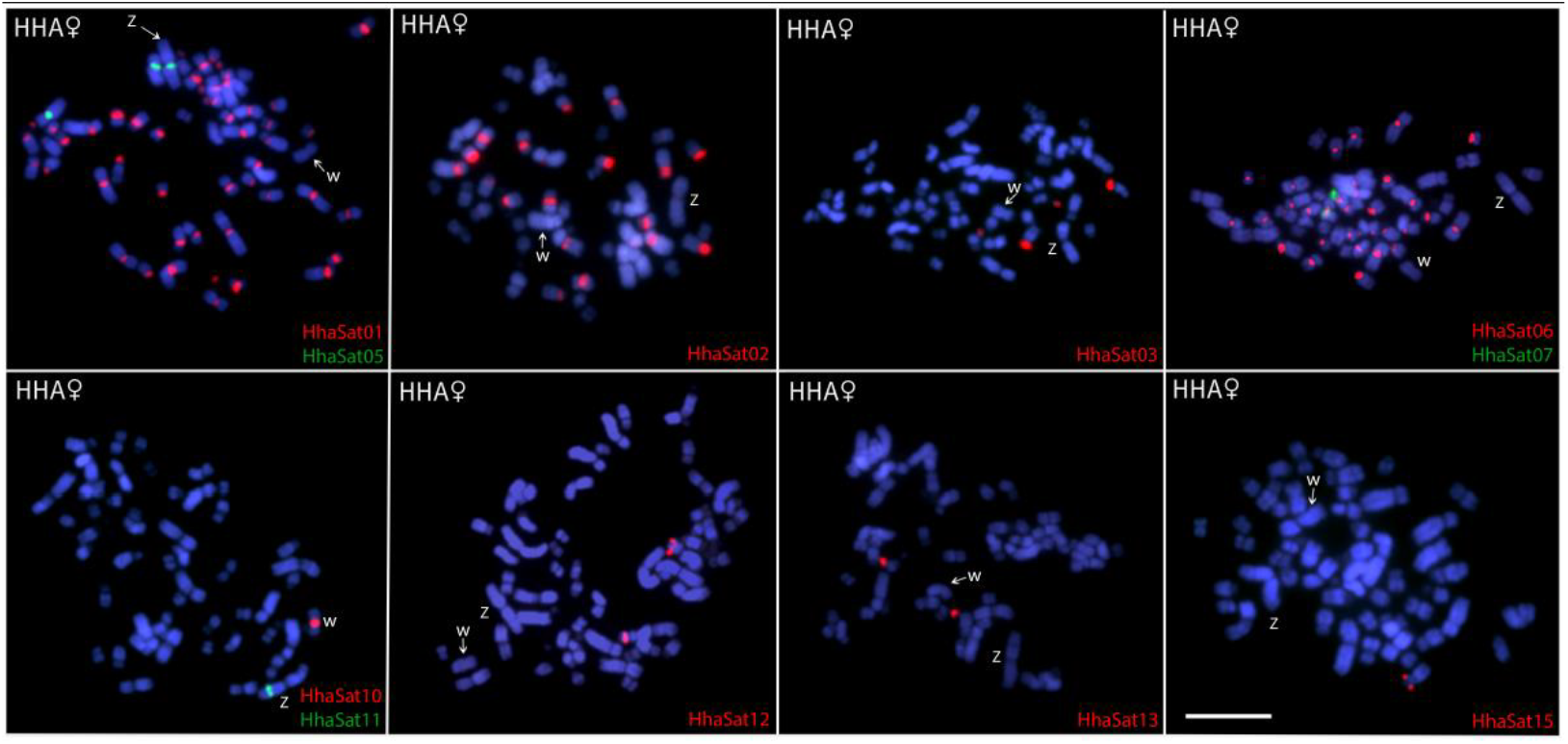
Female metaphasic chromosomes of *Harpia harpyja* showing hybridization signals of HhaSatDNAs as probes. The names of each satellite DNA family are detailed in the bottom-right corner in red (Atto-550-dUTP labeled) or green (ATTO-488-dUTP labeled). The ZW sex chromosomes are indicated. Scale bar= 20 µm.

## Discussion

This study provided an extensive analysis of the evolution of repetitive DNAs in Accipitridae by correlating genome-wide repeat content with their chromosomal locations, thus building a multi-scale framework of genome architecture. Through the integration of comparative repeatome investigations with *in silico* and cytogenetic mapping, we advance from merely establishing repeat abundance to elucidating the spatial organization of repetitive elements across chromosomes. This dual genomic-chromosomal framework enhances the comprehension of repeat dynamics within their structural context, uncovering patterns of conservation, lineage-specific amplification, and spatial clustering that would be bypassed in purely sequence-based approaches, thereby reinforcing the view that repeat landscapes are integrated within structural genomic architecture. This spatial perspective is especially relevant in birds, where long-standing karyotypic stability coexists with lineage-specific bursts of chromosomal reorganization^9,10,12^. In this context, Accipitridae represent a compelling model in which repeat turnover appears tightly coupled to chromosomal restructuring, offering a systems-level view of genome diversification.

The comparative findings herein illustrate that repetitive DNAs are an important driver in genomic architectural variation within Accipitridae and likely influence lineage-specific evolutionary paths. The extensive sharing of SatDNA families among accipitrid species is also consistent with the hypothesis of a shared SatDNA repertoire, which proposes that related taxa inherit a common ancestral set of repeats that is differentially amplified over evolutionary time^28,29^. In our dataset, most SatDNA families belong to conserved groups (superfamilies) present across multiple species **(Supplementary Table 9)**, often retaining similar repeat unit structure and chromosomal distribution, indicating long-term persistence of ancestral elements rather than independent origins. This pattern is particularly evident in Group 2 (**Table 1**) - HhaSat01-173, AguSat04-173, AgeSat01-173, AchSat01-173, BbuSat01-173, and HalSat01-173 - which represents the only SatDNA variant detected across all analyzed species. In addition to its presence in every taxon, this family consistently ranks among the most abundant repeats in each genome and displays relatively high interspecific sequence identity (78.5–93.1%). Taken together, these features support not only an ancestral origin but also long-term evolutionary persistence, suggesting that this repeat family has retained functional or structural relevance throughout the diversification of Accipitridae, as illustrated by the centromeric FISH pattern of HhaSat01 **(Figure 4)**.

Both *in silico* and *in situ* mapping consistently indicate that these sequences are associated with the centromeric regions of most chromosomes, supporting their classification as centromeric SatDNAs **(Figures 3 and 4)**. Previous studies have demonstrated that the most abundant SatDNA families within a genome often correspond to those associated with the functional centromeric domain (centromere-associated), as reported in several avian groups^30-33^. In this context, the high genomic abundance of Group 2, together with the marked structural conservation of its ∼173 bp monomer and its recurrent chromosomal distribution, strongly supports its classification as a centromere-associated SatDNA family **(Table 1)**. In contrast to many mammals, in which alpha satellite DNA constitutes a widely conserved centromeric component across large clades, no universal or broadly conserved centromeric sequence has yet been identified across avian orders^34,35^. Instead, cytogenomic evidence indicates that avian centromeres are highly heterogeneous in sequence composition, even among closely related taxa, reflecting an evolutionary dynamic characterized by rapid satellite turnover and structural reorganization^35,36^. Within this broader context, the pattern observed in Accipitridae is particularly striking: the presence of a shared, highly abundant, and presumably centromeric SatDNA across all analyzed species suggests an unusual degree of centromeric conservation within the lineage. At the same time, species-specific differences in abundance and chromosomal clustering **(Figure 4 and Supplementary Figure 4)** reveal dynamic quantitative turnover, a hallmark of library evolution in which repeat families are maintained but reshaped through lineage-specific amplification and contraction.

In the eight species analyzed, their repeat landscapes exhibit quantitative divergence despite a common structural framework **(Figure 1; Figures Supplementary 2 and 3)**, suggesting that repeat amplification is not random but shifted by lineage. The prevalence of retroelements **(Supplementary Table 2)**, along with species-specific spikes in LINE and LTR expansion **(Figure 1)**, supports a scenario in which episodic repeat proliferation alters genomic structure following a lineage split. Temporal insertion profiles indicate that repeat turnover is an ongoing evolutionary process rather than a relic of ancestral genomic states, aligning with patterns observed in dynamic avian lineages that deviate from canonical genome stability^10,12^. Comparable lineage-specific repeat expansions have been reported in other avian and vertebrate groups, where bursts of transposable element activity correlate with genome restructuring and heterochromatin remodeling^8,11,37^. Notably, species exhibiting marked chromosomal reduction or rearrangement showed enrichment of relatively recent repeats **(Figure 1)**, suggesting that active repeat landscapes may facilitate or accompany structural genome remodeling. Bird genomes are known to have a small portion of repetitive DNAs, around 7-9%^8^, with a few species exceeding this pattern, such as Piciformes^38^ and Strigiformes^39^ with > 20% and sparrow songbirds Passeriformes (Oscines) with > 30%^40^. In accipitrids, species with the greatest deviations from the ancestral avian karyotype exhibit the highest levels of repetitive DNA content **(Figure 1)**, linking repeat expansion to chromosomal remodeling. Elevated repeat fractions across genera further suggest that repeat-rich genomes are a shared feature of this lineage and may underlie its pronounced karyotypic plasticity. Such associations align with broader evidence that repetitive DNA can modulate recombination environments, centromeric function, and chromosome pairing^11,41-43^, thereby influencing karyotypic evolution and potentially speciation trajectories.

Comparative analyses anchored to conserved avian reference genomes (particularly *Gallus gallus* and *D. novaehollandiae*) indicate that breakpoints associated with both inter- and intrachromosomal rearrangements frequently coincide with regions enriched in repetitive DNAs, suggesting that repeat-dense segments may function as intrinsic hotspots of chromosomal instability during avian genome evolution^11,44-46^. Such regions are especially susceptible to ectopic recombination because highly similar repeat sequences are dispersed across nonhomologous chromosomal sites, increasing the likelihood of non-allelic pairing events. These recombination dynamics provide a mechanistic pathway for structural rearrangements (including fusions, fissions, inversions, and translocations) that contribute to karyotypic diversification^47,48^. The high frequency of chromosomal rearrangements documented in Accipitriformes has been interpreted as evidence of a genomic architecture unusually permissive to structural reorganization^14,49^. This karyotypic dynamism differs from patterns seen in other avian lineages and indicates that inherent genomic characteristics may render accipitrid chromosomes susceptible to continual restructuring. In Psittaciformes, for example, extensive chromosomal reconfiguration has been associated with lineage-specific loss of genes involved in double-strand break repair, most notably *ALC1* and *PARP3*, providing a mechanistic link between genome maintenance pathways and structural instability^50^. In contrast, the molecular drivers of rearrangement in Accipitridae remain unresolved, highlighting a critical gap in our understanding of how genome architecture, repair pathways, and repetitive DNA dynamics interact to shape avian karyotype evolution.

Future research should expand sampling across additional accipitrid clades, incorporate long-read and haplotype-resolved assemblies to improve repeat resolution, and combine comparative cytogenomics with functional assays to test how repeat expansions influence chromosomal behavior and gene regulation.

## Material and Methods

### Species

Here, we examined eight Accipitridae species for which reference genome assemblies at the chromosomal level were available **(Table 2)**. Furthermore, a female specimen of *Harpia harpyja* was collected from Mangal das Garças (Belém, Pará) for cytogenetic analysis, as detailed below. The sampling and all technical procedures were conducted according to the Brazilian Environmental Agency ICMBIO/SISBIO license (100206-1).

**Table 2.** Species included in this study, with their diploid chromosome number (2n) and corresponding NCBI accession codes. An asterisk (*) indicates the estimated 2n for *Accipiter gularis* according to Hi-C scaffolding without cytogenetic evidence.

| Species | 2n | Reference | NCBI Assembly accession |
| --- | --- | --- | --- |
| *Buteo buteo* | 68 | De Boer, 1976 | GCA_964188355.1 |
| *Haliaeetus albicilla* | 66 | De Boer, 1976 | GCA_947461875.1 |
| *Astur gentilis* | 78 | De Boer, 1976 | GCF_929443795.1 |
| *Accipiter gularis* | 66* | - | GCA_046127185.1 |
| *Morphnus guianensis* | 54 | Williams & Benirschke, 1976 | GCA_045345505.1 |
| *Harpia harpyja* | 58 | De Oliveira et al. 2005 | GCF_026419915.1 |
| *Aquila chrysaetos* | 62 | Takagi & Sasaki, 1974 | GCF_900496995.4 |
| *Gypaetus barbatus* | 60 | De Boer, 1976 | GCA_028022735.1 |

### Chromosomal Preparation, DNA extraction, and Genome sequencing of H. harpyja

A fibroblast cell culture was established using skin biopsies from a female *H. harpyja*, and mitotic chromosomes were obtained following the protocol described by^51^. Genomic DNA was extracted from the cell culture according to the protocol described by^52^. All experiments were conducted in strict accordance with the ethical guidelines and regulations of the Animal Experimentation Ethics Committee of the Universidade Federal de São Carlos (CEUA-7994170423). Paired-end libraries with a read length of 150 bp were generated from each genomic DNA, followed by low-coverage sequencing conducted on the BGISEQ-500 platform at BGI (BGI Shenzhen Corporation, Shenzhen, China). The sequencing produced a total of 3 Gb of raw reads, corresponding to ∼2X genomic coverage.

### Repeatome characterization

The repeatome analysis included the data generated here for *Harpia harpyja* (as indicated above) and short-read sequencing for additional species retrieved from public databases (NCBI/GenBank): *Aquila chrysaetos* (DRX346971), *Astur gentilis* (ERX14695606), *Accipiter gularis* (SRX26100943), *Buteo buteo* (SRX15141501), and *Haliaeetus albicilla* (ERR16057181). Satellite DNA sequences from short-read data were characterized using the RepeatExplorer and TAREAN pipelines^21,22^, implemented on the Galaxy server (https://repeatexplorer-elixir.cerit-sc.cz/galaxy/), following the protocol described by^53^. All putative satellite sequences were subjected to a multi-step manual curation process. First, sequences showing homology to multigene families were removed. Next, homology analyses were performed using the rm_homology script (https://github.com/fjruizruano/satminer/blob/master/rm_homology.py) in combination with the Cross_match search engine implemented in RepeatMasker^54^. Clustered sequences were then aligned and classified based on pairwise identity: sequences sharing >80% identity were considered variants of the same SatDNA family, whereas those sharing >50% identity were grouped into the same superfamily, following^53^. Transposable elements (TEs) were characterized de novo using RepeatModeler2^55^ with the -LTRStruct parameter using the genome assemblies listed in **Table 2**. Satellite DNAs for species lacking short-read data (*G. barbatus* and *M. guianensis*) were identified using the Satellite Repeat Finder (SRF)^56^. Repetitive elements detected with RepeatModeler and satellite sequences identified via TAREAN or SRF were combined into a unified repeat library. This library was then used to estimate genome-wide repeat abundance and sequence divergence with RepeatMasker^54^. Repeat insertion timing was calculated using parseRM (https://github.com/4ureliek/Parsing-RepeatMasker-Outputs), which parses RepeatMasker outputs, applying a neutral mutation rate of 2.3 × 10^−9^ substitutions per site per year^57,58^.

A heatmap of repetitive content for each species was generated using chromosome-level genome assemblies **(Table 2)**. The analysis was based on the RepeatMasker .gff output files^54^, and repetitive elements were quantified in non-overlapping 50-kb windows to improve the visualization of abundance patterns across specific genomic regions. The R package RIdeogram^59^ was used to visualize the density of repetitive elements along the chromosomes.

### Primer Design and Probe Construction

Primers were designed exclusively to amplify SatDNAs identified in *Harpia harpyja* for *in situ* chromosome mapping **(Supplementary Table 1)**. Only SatDNA families with sufficient sequence length to support a reliable primer design were considered. Two satellites (HhaSat08 and HhaSat09) were excluded due to high sequence homology with other satellite families (see **Table 1**). The shortest satellites (<35 bp; HhaSat02, HhaSat03, and HhaSat13) were synthesized directly, incorporating biotin molecules at the 5′ end. The amplification protocol consisted of an initial denaturation step at 95 °C for 5 minutes, followed by 32 cycles of 95 °C for 20 seconds, annealing at 54–60 °C for 40 seconds, 72 °C for 50 seconds, and a final extension step at 72 °C for 5 minutes. Agarose gel electrophoresis (2%) was then used to confirm amplification, and the resulting products were quantified using a NanoDrop spectrophotometer (ThermoFisher Scientific, Branchburg, NJ, USA). PCR products were labeled by nick translation for probe construction and subsequently used in *in situ* hybridizations, employing Atto488-dUTP and Atto550-dUTP nucleotides (Jena Biosciences, Jena, Germany).

### In situ Mapping

Fluorescence *in situ* hybridization (FISH) experiments were conducted as established by^60^. The mixture used for hybridization included 200 ng of labeled probe, 50% formamide, 2× SSC, 10% SDS, and 10% dextran sulfate (Denhardt’s buffer, pH 7.0), all adding up to a total of 20 µl.

### Statistics and reproducibility

At least 20 metaphases were examined to confirm the diploid number (2n) and the FISH results. The best metaphases were captured using an Olympus BX50 microscope (Olympus Corporation, Ishikawa, Japan) coupled with the CoolSNAP capture system and Image Pro Plus 4.1 software (Media Cybernetics, Silver Spring, MD, USA).

### In silico Mapping of SatDNAs

For *in silico* mapping, we used SatDNA families identified as variants **(Table 1)**. In species lacking independently characterized satellitomes (*G. barbatus* and *M. guianensis*), mapping was performed using *the H. harpyja (HhaSatDNA) and B. buteo (BbuSatDNA) satellite families*, both belonging to shared satellite groups. Satellite sequences were aligned against chromosome-level genome assemblies of Accipitridae species **(Table 2)** using BLASTn^61^. Alignments were filtered using stringent quality thresholds, including e-value, percentage identity, and minimum alignment length, to exclude low-confidence matches. Only filtered hits were retained for downstream analyses. Genomic coordinates were exported in BED format and visualized along chromosomes. SatDNA visualization on chromosomes was performed using the karyoploteR package in R^62^, which enables the construction of chromosomal ideograms and the mapping of sequences along their lengths. To create the plots, in addition to the “.bed” files containing filtered hits, a “.genome” file was used, containing one column with chromosome names and another with size in base pairs, enabling the analysis of location, relative abundance, and distribution patterns of satellite sequences across chromosomes of different species.

### Synteny analysis

For synteny analysis, we used the NUCmer pipeline^63,64^. NUCmer operates solely on DNA sequences, directly comparing genomic sequences and identifying regions of nucleotide similarity and collinearity. In addition to the selected Accipitridae species, the genomes of the king vulture (*Sarcoramphus papa*) and emu (*Dromaius novaehollandiae* (genome accessions: GCA_037962945.1 and GCA_036370855.1, respectively) were included in the comparative analyses as outgroups. The NUCmer pipeline involved five main steps: (i) identification of collinear regions between the two sequences using the nucmer command (requiring a single file for each species containing all chromosomes), (ii) filtering the alignments with delta-filter, (iii) converting the filtered .delta file into a plottable format with show-coords, (iv) converting the .coords files to .bed and .simple formats, and (v) plotting the results using the MCScan program from the JCVI package^65^. The final images were manually refined and formatted in Adobe Illustrator 2020 (v. 24.1.2).

## Data availability

Sequencing data that support the findings of this study have been deposited in the Sequence Read Archive (SRA-NCBI) under accession number PRJNA1427246, corresponding to the raw genomic data of *Harpia harpyja*. Satellite DNA (SatDNA) sequences characterized in this study have been deposited in the GenBank database under the following accession numbers: PZ005776–PZ005790 (*Harpia harpyja*), PZ005791–PZ005806 (*Buteo buteo*), PZ005807–PZ005814 (*Haliaeetus albicilla*), PZ005815–PZ005832 (*Aquila chrysaetos*), PZ005833–PZ005850 (*Accipiter gularis*), and PZ005851–PZ005866 (*Astur gentilis*). These accession numbers provide public access to all SatDNA consensus sequences analyzed in the present study.

## Acknowledgments

Authors were supported by the Fundação de Amparo à Pesquisa do Estado de São Paulo (FAPESP) for G.M.S. (Proc. 2025/10370-7) and M.B.C. (Proc. 2024/12644-4). E.H.C.d.O. (Proc 307382/2019-2) and M.B.C. (Proc. 302928/2021-9) were also supported by the Brazilian National Council for Scientific and Technological Development (CNPq). We also thank Bruno Francelino de Melo for his assistance with the adjustments in the phylogeny presented in Figure 1. This study was supported by the Coordenação de Aperfeiçoamento de Pessoal de Nível Superior (CAPES, finance code 001). We also acknowledge support by the German Research Foundation Projekt-Nr. 512648189 (T.L.) and the Open Access Publication Fund of the Thueringer Universitaets- und Landesbibliothek Jena. For open access purposes, the authors have applied the Creative Commons CC BY license to any accepted version of the article.

## Author contributions

G.M.S., R.K. and M.B.C. conceived and designed the research. G.M.S., J.A.D.V., G.A.T. and E.H.C.O. conducted the experiments. G.M.S., J.A.D.V., G.A.T., R.K., E.H.C.O., T.L. and M.B.C analyzed the data. G.M.S., J.A.D.V. and G.A.T. contributed with new methods. G.M.S., J.A.D.V., G.A.T., R.K., E.H.C.O., T.L. and M.B.C wrote the paper

## Competing interests

The authors declare no competing interests.

## Additional information

### Supplementary information

The online version contains supplementary material

## Notes

### Competing Interest Statement

The authors have declared no competing interest.

