## Supplementary material for "Repetitive DNA shapes genome architecture and chromosomal diversification in birds of prey": Supplementary Files.docx

**Supplementary Figures**

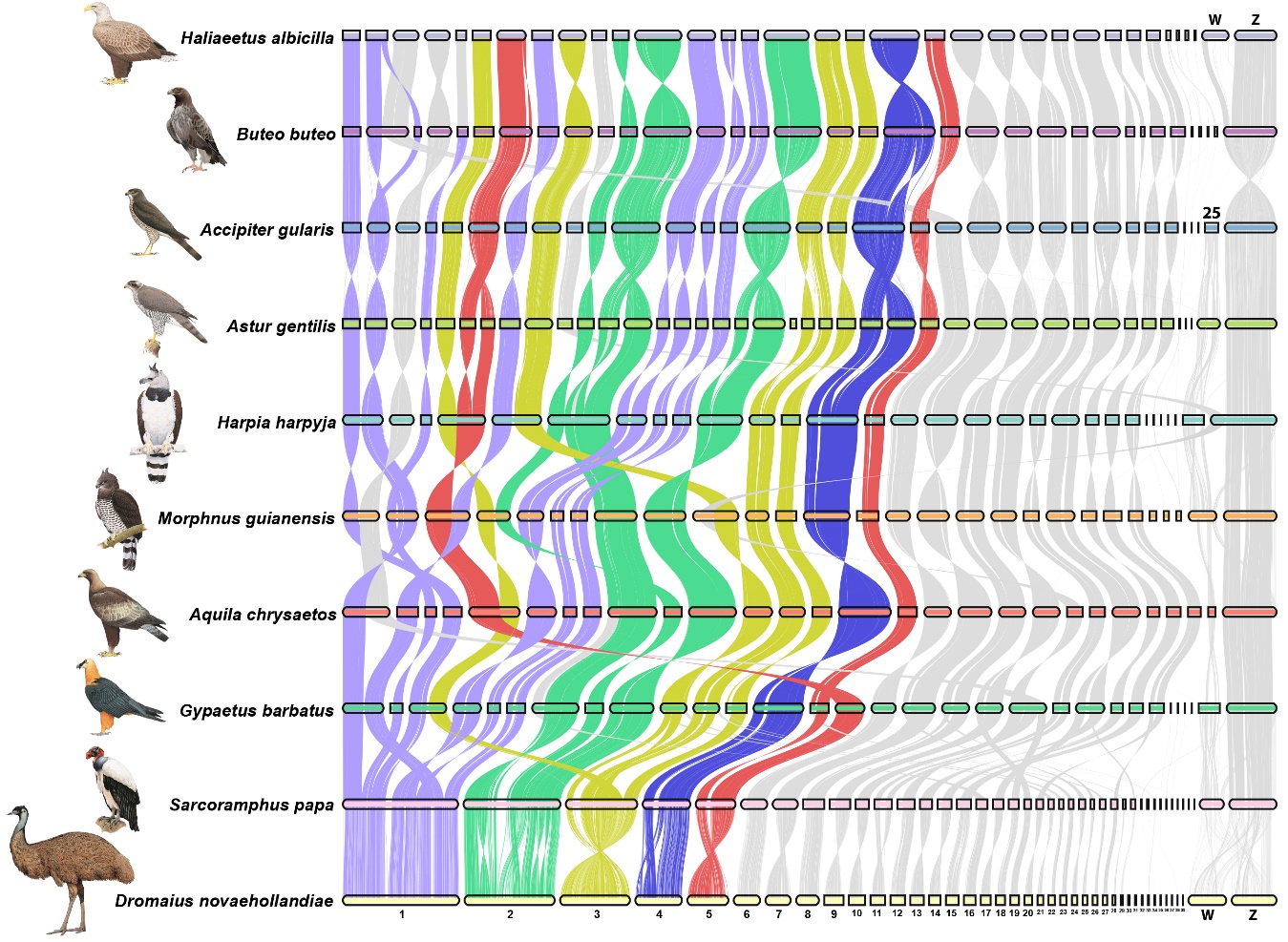

**Supplementary Figure 1.** Pairwise whole-genome alignments across eight chromosome-level assemblies of Accipitriform genomes. The species *Sarcoramphus papa* (Cathartiformes) and *Dromaius novaehollandiae* (Casuariiformes) were included as outgroups. Each horizontal bar represents a chromosome, and chromosomes that underwent recurrent chromosomal rearrangements are highlighted in color.

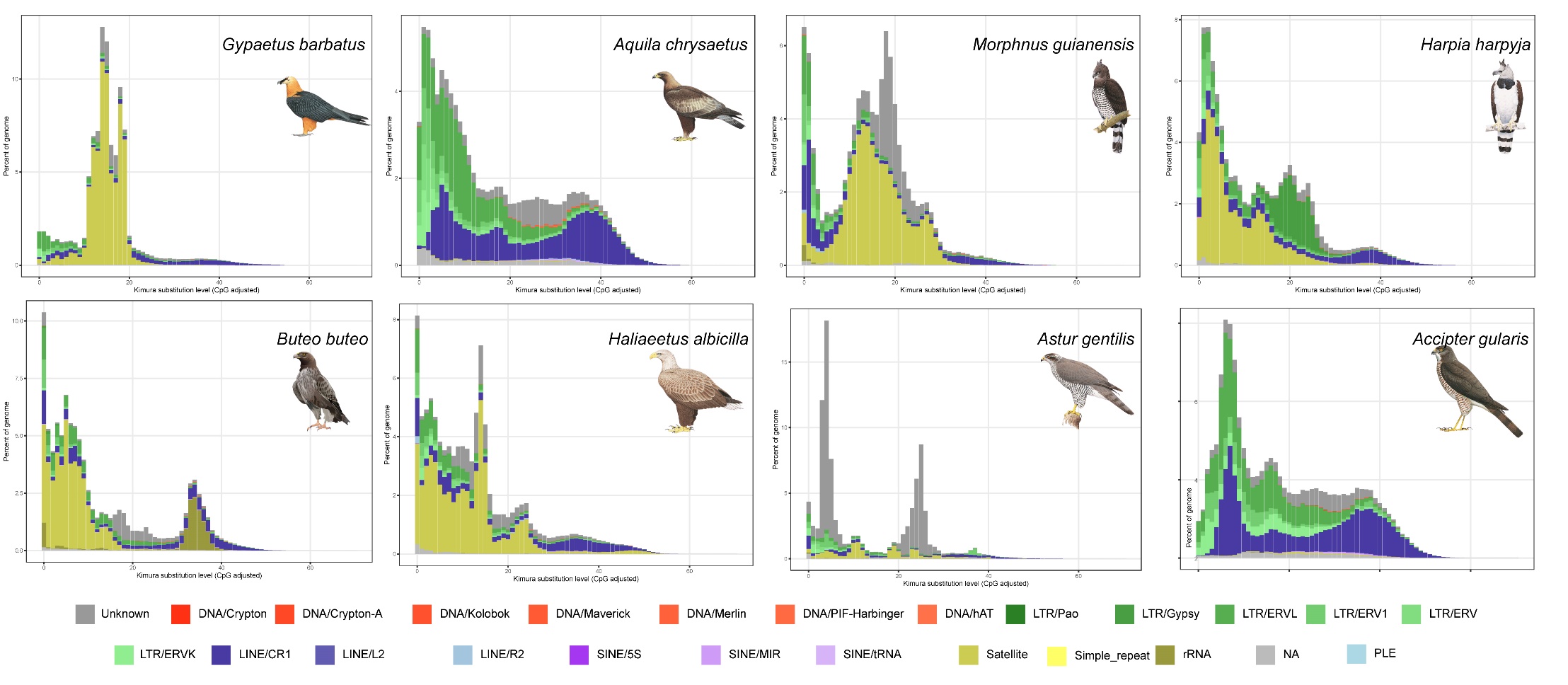

**Supplementary Figure 2.** Repeat landscapes showing the abundance and divergence profiles for all repetitive DNAs identified in *Aquila chrysaetos*; *Harpia harpyja; Astur gentilis; Accipiter gularis; Buteo buteo*, and *Haliaeetus albicilla*.

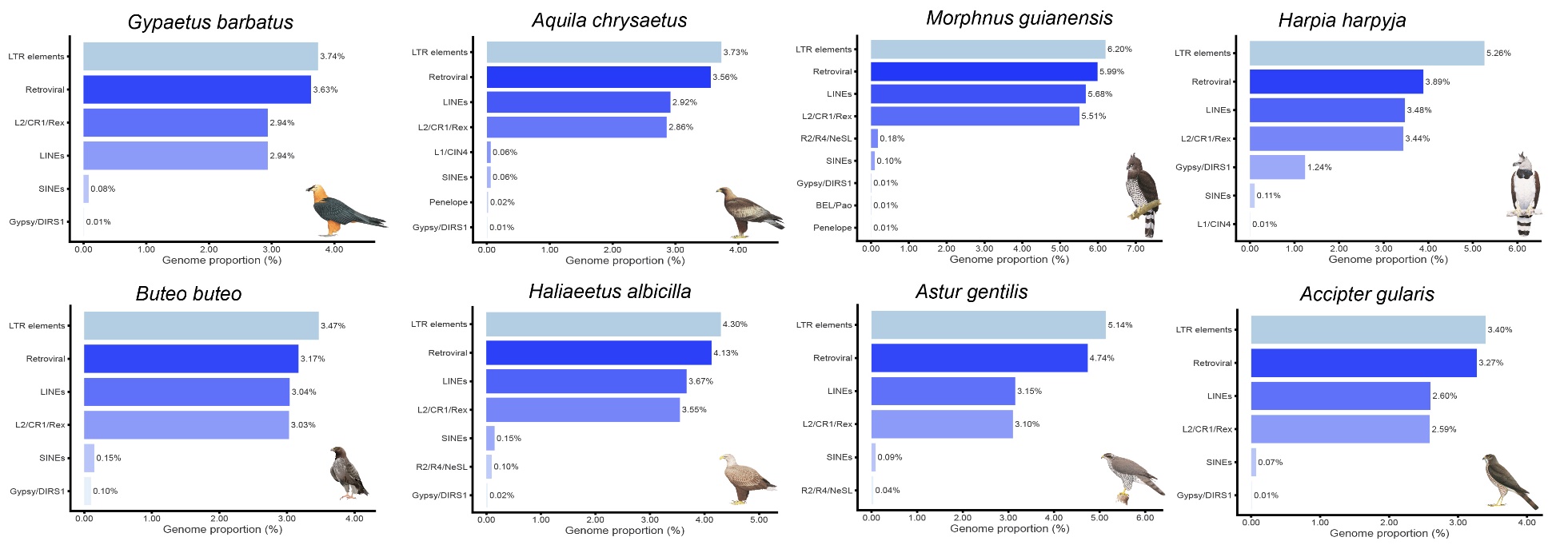

**Supplementary Figure 3.** Genome proportion of the main retroelements present in the genomes of the eight Accipitridae species analyzed in this study.

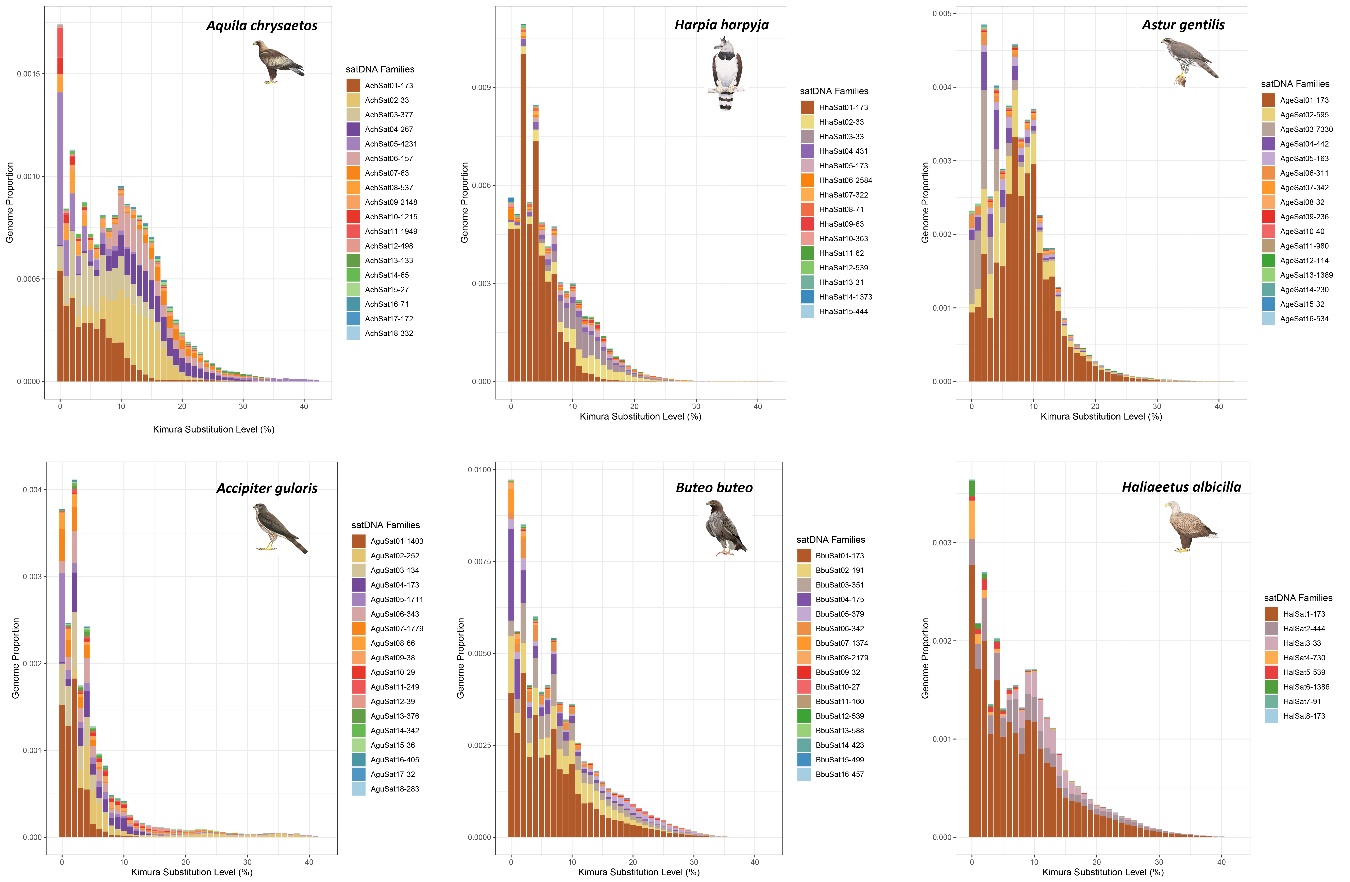

**Supplementary Figure 4:** Repeat landscapes showing the abundance and divergence profiles for all satDNAs identified in *Aquila chrysaetos*; *Harpia harpyja; Astur gentilis; Accipiter gularis; Buteo buteo*, and *Haliaeetus albicilla*.
